# Scalable bio-platform to recover critical metals from complex waste sources

**DOI:** 10.1101/2023.09.02.556034

**Authors:** Nathan M. Good, Christina S. Kang-Yun, Morgan Z. Su, Alexa M. Zytnick, Huong N. Vu, Joseph M. Grace, Hoang H. Nguyen, Dan M. Park, Elizabeth Skovran, Maohong Fan, N. Cecilia Martinez-Gomez

**Affiliations:** Department of Plant and Microbial Biology, University of California, Berkeley, Berkeley, California, USA; Physical and Life Sciences Directorate, Lawrence Livermore National Laboratory, Livermore, California, USA; Department of Biological Sciences, San José State University, San José, California, USA; Department of Chemical and Biomedical Engineering, University of Wyoming, Wyoming, USA

**Keywords:** bioaccumulation, bioconcentration, electronic waste, lanthanide, metal-binding protein, neodymium, bioleaching, acid-free leaching

## Abstract

Chemical methods for extraction and refinement of technologically critical rare earth elements (REEs) are energy intensive, hazardous, and environmentally destructive. Current bio-based extraction systems rely on extremophilic organisms and generate many of the same detrimental effects as chemical methodologies. The mesophilic methylotrophic bacterium *Methylobacterium extorquens* AM1 was previously shown to grow using electronic waste by naturally acquiring REEs to power methanol metabolism. Here we show that growth using electronic waste as a sole REE source is scalable up to 10 L with consistent metal yields, without the use of harsh acids or high temperatures. Addition of organic acids increases REE leaching in a nonspecific manner. REE-specific bioleaching can be engineered through the overproduction of REE-binding ligands (called lanthanophores) and pyrroloquinoline quinone. REE bioaccumulation increases with leachate concentration and is highly specific. REEs are stored intracellularly in polyphosphate granules, and genetic engineering to eliminate exopolyphosphatase activity increases metal accumulation, confirming the link between phosphate metabolism and biological REE use. Finally, we report the innate ability of *M. extorquens* to grow using other complex REE sources including pulverized smart phones, demonstrating the flexibility and potential for use as a recovery platform for these critical metals.

## INTRODUCTION

Global demand for rare earth elements (REEs) is at an all-time high and is steadily increasing, but their supply is susceptible to market and national interest perturbations.[1–3] REEs, composed of the lanthanides (Lns), scandium, and yttrium, are critical metals for modern clean energy, communication, advanced transportation, consumer electronics, and defense technologies, underpinning a broader economy.[4–6] China maintains its status as the world’s foremost producer of REEs, generating heavy reliance on a single market that could jeopardize supply and compromise national security.[7,8] Though domestic REE production is on the rise, environmental and health concerns loom over traditional mining operations.[9],[10,11] The development of technologies for REE reuse and recycling has garnered interest as a means of moving towards independence from foreign importation while simultaneously generating a robust, resilient supply of these critical metals.[12]

Several challenges continue to stymy development of a safer, more environmentally conscious, REE supply chain. These include, (1) the requirement of high temperatures and pressures, and inclusion of harsh acids for extraction of poorly soluble REE, which can produce radioactive waste products; (2) the preference for high-grade sources (>0.2% REE m/m) for cost-efficient recovery, leaving low-grade and waste sources as untapped reservoirs of valuable REE; and (3) the necessity for hundreds of costly, hazardous processing steps for successful separation of co-occurring REE in minerals.[2,13]

Microbiological REE leaching and extraction methods offer promising alternatives to current state-of-the-art methods (hydrometallurgical, pyrometallurgical, and electrometallurgical approaches) that produce large quantities of sludge, acidic wastewater, atmospheric pollution, and radioactive tailings.[9,14–16] Given that conventional rare earth ores (e.g., bastnaesite, monazite, xenotime) are typically oxidized and non-sulfidic[17], conventional bioleaching using acidophilic sulfur and iron oxidizers, which has been widely practiced commercially for sulfidic copper and gold ores, is not directly applicable. Rather, REE bioleaching approaches typically involve the production of organic acids (e.g., citric, gluconic, oxalic acids)[18,19] from sugar-based carbon sources by heterotrophic bacteria or fungi to leach REEs into solution. While these approaches can be effective for overall leaching, and even provide an economical benefit,[19–21] they indiscriminately dissolve metal feedstocks and require an additional process(es) to separate desired metals from the remaining leachate. Microbial-mediated REE adsorption using native and engineered cell surface functional groups (e.g., carboxylates, phosphates, and metal binding outer membrane proteins) [22–25] can be an effective and potentially economic strategy [26,27] for selective sequestration of REE from bulk samples, but this method requires solution-based feedstocks derived from costly chemical leaching and pH adjustment processes.

Microbial metal accumulation and biomineralization has been shown to be an effective strategy for recovery of Hg,[28] Au,[29–31] and U,[32,33] but has been underexplored as an REE recovery approach. Bioaccumulation and biomineralization of REEs in mesophilic bacteria was first shown in the model methylotroph *Methylobacterium* (also known as *Methylorubrum*) *extorquens* AM1.[34] Methylotrophic bacteria, organisms that thrive on inexpensive, readily available one-carbon compounds such as methane and methanol, therefore provide an attractive new approach for REE bioleaching and extraction due to their natural ability to acquire Lns from the surrounding environment.[35,36] This includes soluble and insoluble REE sources, such as electronic waste (E-waste).[37–39] *M. extorquens* AM1 was the first organism reported to grow using REE from electronic waste.[36] Mesophilic methylotrophs like *M. extorquens* AM1 have dedicated systems for acquisition, uptake, and intracellular storage of Lns as polyphosphate granules, making them effective agents of bioleaching and bioaccumulation without the need for high acidity or temperature.[34] REE use by mesophilic methylotrophs was thought to be restricted to the light Ln, but recently, a genetic variant of *M. extorquens* AM1 was isolated and characterized that can transport, store, and grow using the heavy Ln, gadolinium.[40] Detailed genetic and Ln uptake studies indicate the likely possibility of an additional system dedicated to acquisition and transport of heavy Lns. Thus, methylotrophs, such as *M. extorquens* AM1, may already possess the biological means to separate light and heavy Lns, and have the potential to be engineered for uptake of specific Ln species from complex feedstocks. To date, the potential of methylotrophs as a platform for REE recovery has not been rigorously investigated, including the selectivity of REE uptake.

Here, we show the scalability of this growth up to 10 L with Nd magnet swarf, waste powders and filings generated during magnet production and an E-waste analog without the requirements of high temperatures or harsh acids. We also show the potential for process improvement by leveraging the genetic tractability and suite of genetic tools available for *M. extorquens* AM1 to engineer enhanced non-acidic REE bioleaching (∼20-fold) and bioaccumulation (∼50-fold) attributes. This study provides a proof-of-principle demonstration of *M. extorquens* AM1 as a scalable platform for biomining and bioaccumulation of REEs from complex feedstocks, such as magnet swarf, without the need of harsh acids and high temperatures, greatly reducing potential hazards and obstacles to generating a circular economy (Figure 1).

**Figure 1.**
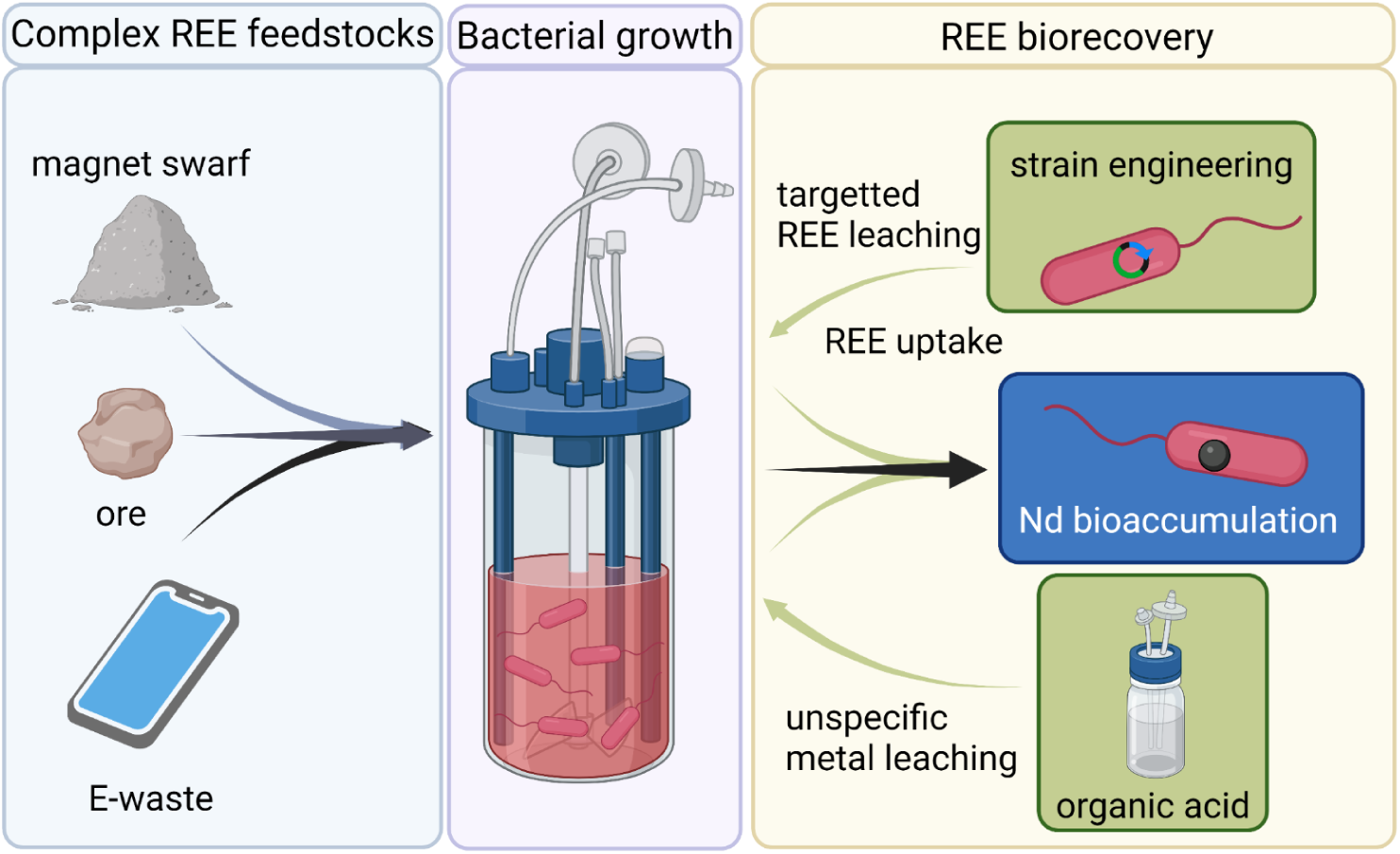
*M. extorquens* AM1 is a platform for Nd recovery through efficient, high-yield bioaccumulation from complex sources. REEs in complex feedstocks are accessible to *M. extorquens* AM1 due to its innate ability to selectively acquire, transport, and accumulate the metals in polyphosphate granules. Metal leaching and REE uptake can be enhanced by addition of organic acid to the culture medium and strain engineering, resulting in greater Nd recovery yields.

## MATERIALS AND METHODS

### Chemicals

All chemicals were purchased through Fisher Scientific (Pittsburgh, PA) or Sigma-Aldrich (St. Louis, MO) unless otherwise specified. Mineral ores containing lanthanides were purchased from vendors via Ebay.com. Magnets were removed from their protective stainless-steel case prior to processing. Lanthanide-containing ores and magnets were powderized for subsequent experiments. NdFeB magnet swarf was provided by Urban Mining Company (Austin, TX).

### Bacterial strains and growth conditions

All growth studies were performed using the Δ*mxaF* strain of *M. extorquens* AM1 to ensure REE dependency for growth.[41] For routine culturing of *M. extorquens* AM1, strains were grown in 16 x 125 mM borosilicate culture tubes shaking at 200-220 rpm, 29-30°C in modified *Methylobacterium* PIPES (MP) or Hypho minimal medium containing 2-20 µM calcium chloride,[41,42] with 1.75 g/L succinate as a sole carbon source. Except when indicated otherwise, a modified formulation of Hypho (Hypho*^MOD^*) containing 1.27 g/L K_2_HPO_4_ and 1.30 g/L NaH_2_PO_4_.H_2_O was used. For growth measurements with solid ores and minerals, cells were grown in 16 x 125 mm or 25 x 150 mm borosilicate culture tubes using an Excella E25 shaking incubator (New Brunswick Scientific, Edison, NJ) at 180 rpm.

### Growth analysis with organic acids

*M. extorquens* AM1 Δ*mxaF* was grown overnight in 3 mL of MP medium with 1.77 g/L succinate as a sole carbon source at 30 °C, 220 rpm in an I 24 shaking incubator (New Brunswick Scientific, Edison, NJ). Cells were harvested via centrifugation at 6,900 × g for 10 min (Centrifuge 5420; Eppendorf, Hamburg, Germany), then washed with 1 V of fresh, Hypho*^MOD^* medium. The resulting cell pellet was resuspended in 0.1 V of fresh medium, then used to inoculate 1-mL cultures in 24-well suspension culture plates (Greiner Bio-One, Monroe, NC) at a final dilution rate of 1:10. Control cultures were incubated at 30 °C, 567 rpm in a microplate reader (BioTek Synergy H1; Agilent, Santa Clara, CA), and growth was monitored continuously by measuring optical density at 600 nm (OD_600_). Cultures with magnet swarf were incubated at 30 °C, 500 rpm in a Thermoshaker PHMP-4 (Grant Instruments, Royston, U.K.), and periodically removed to measure OD_600_ in the microplate reader. These cultures were passed over a permanent magnet to clear the well center of magnet swarf particles prior to measurements. To investigate the effect of organic acids on methylotrophic bioleaching and bioaccumulation with 1% (w/v) Nd magnet swarf and 1.6 g/L methanol as sole sources of REE and carbon, respectively, cultures were grown with and without 5 mM or 15 mM citrate, 55 mM gluconate, or 5 mM oxalate. The cells were harvested at early stationary phase by transferring the culture into a 1.5-mL microcentrifuge tube to separate from the remaining magnet swarf. The cultures were then centrifuged, and cell pellet and spent medium fractions were separately stored at −20 °C. Biological triplicates were analyzed for every condition.

### Bioreactor cultivation

Pre-cultures grown in Hypho*^MOD^* medium with succinate were prepared as mentioned and then sub-cultured into 50 mL of Hypho*^MOD^*medium with 1.75 g/L succinate and 1.6 g/L methanol in 250 mL shake flasks. Cultures were incubated shaking at 200 rpm, 30°C in a Innova S44i shaker (Eppendorf, Hamburg, Germany) for 16 hours. Thirty mL of culture was then transferred to a 2 L BIOne 1250 bioreactor (Distek, North Brunswick, New Jersey) with 0.75 L of Hypho*^MOD^*medium and 1% (w/v) NdFeB magnet swarf. For 10 L bioreactor cultivations, 500 mL of overnight cultures was transferred to 9.5 L of Hypho*^MOD^*medium. Bioreactor parameters were as follows: agitation, 500 rpm; air flow, 200-2000 sccm; temperature 29.5°C; pH 6.9. The pH of the bioreactor was maintained using 1 M NaOH.

### Metal quantification

Samples for the determination of Nd content were prepared as follows: 10 mL of culture was removed from the bioreactor and cells were pelleted by centrifugation at 4121 x *g* in a Multifuge X Pro Series centrifuge (Thermo Fisher Scientific, Waltham, MA) for 12 minutes. Culture supernatant was decanted and cell pellets were washed four times with double-distilled water before drying at 65°C. After achieving complete dryness, dry weight was measured, and then cells were deconstructed in 20% metals-grade nitric acid at 90°C and diluted to 2.3% acid before analysis by inductively coupled plasma mass spectrometry (ICP-MS) at the Laboratory for Environmental Analysis (Center of Applied Isotope Studies, University of Georgia). Values were determined by ICP-MS and normalized per unit dry weight as recommended for comparison of bioaccumulation across samples and methods.[43]

Cell pellets obtained from growth in 24-well plates were acidified in 35% (v/v) HNO_3_ for 2 h at 95 °C with mixing by inversion every 30 min. Prior to dilution, the REE concentrations in spent medium and acidified cell samples were quantified by the Arsenazo III assay, as described previously.[44] Briefly, 40 μL of sample was combined with 40 μL of 12.5% (w/v) trichloroacetic acid (TCA), then added with 120 μL of filtered 0.1% (w/v) Arsenazo in 6.25% (w/v) TCA. Absorbance at 652 nm was measured and REE concentration was calculated based on a calibration curve with Nd. For metal content analysis using ICP-MS, digested cell suspensions and spent medium samples were diluted with Milli-Q water or 4% (v/v) HNO_3_, respectively, to achieve a final HNO_3_ concentration of 2%. The amount of metals associated with cells was normalized to biomass using a predetermined factor of 0.422 ± 0.018 mg dry weight mL^-1^ OD_600_^-1^ (n = 5).

### Statistical analysis

Data were analyzed using one-way analysis of variance (ANOVA) followed by Tukey’s honestly significant difference (HSD) test to determine significant differences between conditions. Maximum growth rates in 24-well cultures were determined via linear fitting of natural log transformed OD_600_ using the R package growthrates.[45]

### Molecular cloning and mutagenesis

The Δ*ppx* mutant strain was generated using the counter selection marker *sacB*. The donor plasmid was constructed as follows: ∼800 bp regions of genomic DNA flanking the *ppx* gene were amplified using primers designed with 20 bp overlaps for homology-based assembly as previously reported [46]. Linearized pCM433KanT was also produced by PCR with 20 bp overlaps for the 5’ and 3’ flanking regions. The final construct was assembled as previously described [40], Sanger sequenced for verification, and transformed into *M. extorquens* AM1. Counterselection was conducted with 5% sucrose as reported [47]. The deletion was confirmed via PCR and Sanger sequencing.

## RESULTS AND DISCUSSION

### Scalable REE bioleaching and bioaccumulation from electronic waste

*M. extorquens* AM1 cultures grown in Hypho*^MOD^* (see Methods, Supplemental Methods, Figure S1) in a 0.75 L benchtop bioreactor and 1% (m/v) Nd magnet swarf generated growth rates that were significantly increased compared to microplate (+2.3-fold) and shake flask cultures (+1.5-fold) (Table 1). In the bioreactor, cultures reached maximum densities in 25 h, representing a 2.2-fold reduction in growth period relative to culturing in microplate and a 3.2-fold reduction compared to growth in shake flasks (Table 1). At higher pulp densities, leaching of REEs and/or other metals, such as iron, could negatively impact growth performance and ultimately bioaccumulation. However, when magnet swarf was increased to 10 % (m/v) in the same bioreactor setup, growth rates and cycle times were similar to what was observed at the lower pulp density, indicating that growth was not inhibited.

**Table 1.**
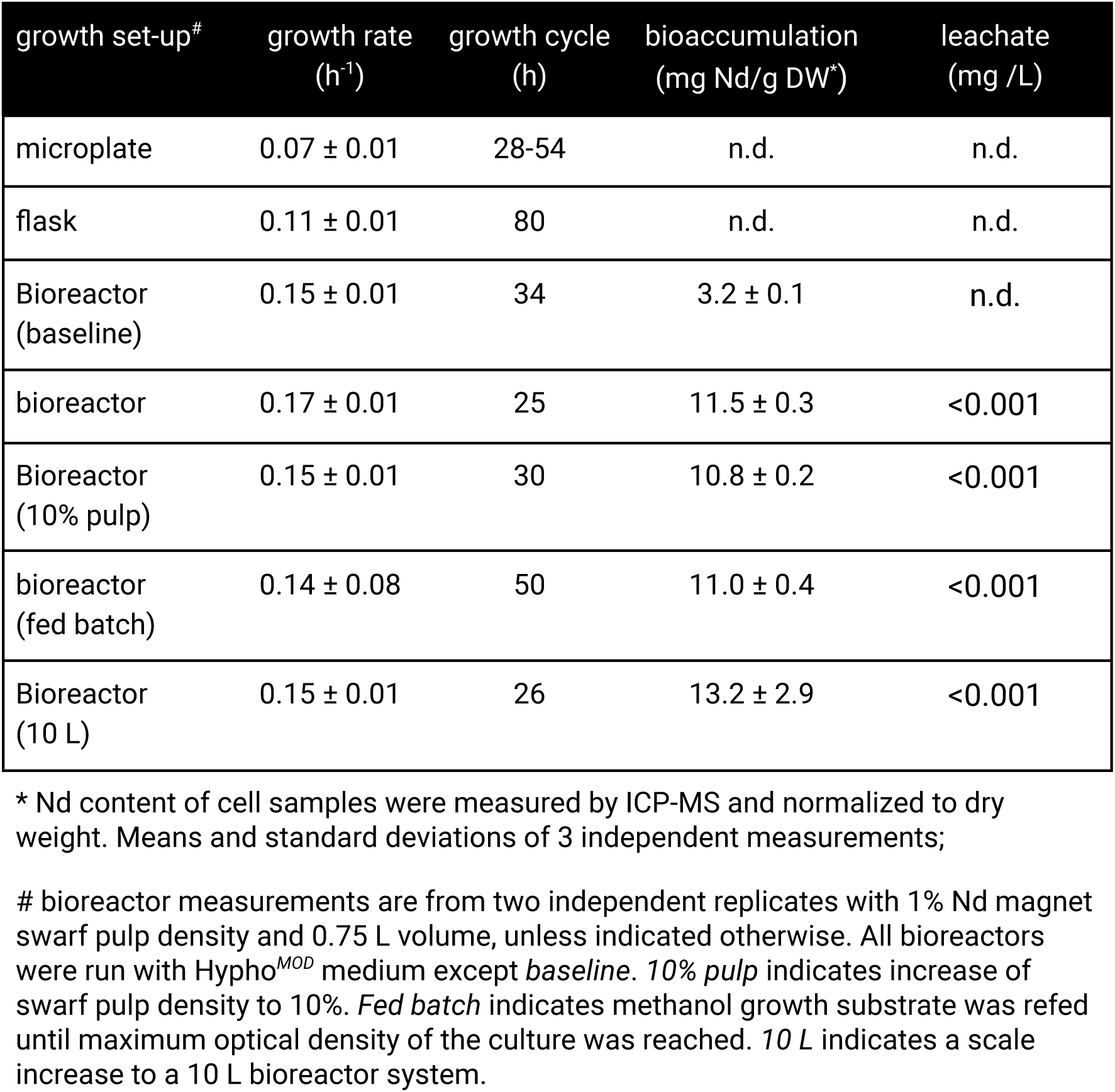
Growth performance and bioaccumulation of Nd by *M. extorquens* AM1 during bioreactor cultivation with Nd magnet swarf.

*M. extorquens* AM1 stores REE as intracellular polyphosphate granules.[34] Baseline Nd bioaccumulation levels were measured from bioreactor cultures grown in standard Hypho minimal medium (Table 1). Relative to the baseline bioaccumulation level, *M. extorquens* AM1 grown in Hypho*^MOD^* accumulated 3.6-fold more Nd when grown with 1% pulp density (Table 1). Reducing inorganic phosphate, therefore, not only boosted growth performance in the bioreactor, but also positively affected REE bioaccumulation, underscoring the link between phosphate metabolism and REE use in microbial systems. Removal of calcium from the growth medium did not impact REE bioaccumulation (Figure S2).

State-of-the-art genetically engineered microbes for heavy metal bioaccumulation have achieved recoveries of up to a few mg/g dry weight, though there are some notable exceptions, such as copper, with bioaccumulation approaching 150 mg/g dry weight.[43] In addition, most studies have reported bioaccumulation from starting concentrations of < 75 mg heavy metal/L. A pulp density of 1% magnet swarf is the equivalent of ∼2 g Nd/L and ∼5.5 g Fe/L if all the magnet was dissolved. Even at 10% pulp density, REE bioaccumulation by *M. extorquens* AM1 was still high (Table 1) indicating that the bacterium is naturally tolerant to high concentrations of heavy metals. Astonishingly, we detected very little residual Nd in the leachate in either growth condition (Table 1) showing that *M. extorquens* AM1 is naturally adept at solubilizing and acquiring REE from complex sources like Nd magnet swarf.

Next, we tested the performance of *M. extorquens* AM1 in methanol fed-batch conditions in a 0.75 L bioreactor. By repeated feeding of methanol over the run cycle, culture densities of 20 OD were achieved without a significant reduction in growth rate (Table 1). In total, ∼17.3 g/L methanol was fed over a cycle time of ∼50 hours. Again there was no detectable residual Nd found in the leachate and the REE to biomass ratio was the same as in batch cultures (Table 1). Finally, we assessed process performance at 10 L scale using optimized media and 1% Nd magnet swarf pulp density. Both growth cycle and growth rate were like the 0.75 L scale, and Nd yields increased by nearly 15%. Together, these results show the scalability of this process for efficient REE recovery from E-waste.

### Selective bioaccumulation of REE from Nd magnet swarf

Even high-grade feedstocks like Nd magnet swarf are of mixed composition, and therefore vary in the content of highly valuable REE and associated metals,[48] making selectivity crucial for the development of a successful recovery platform. We determined the precise metal content of the Nd magnet swarf used in our growth and recovery experiments using ICP-MS. Iron was the primary metal component of Nd magnet swarf (68.0% Fe), rendering non-selective leaching and uptake mechanisms insufficient for effective REE bioaccumulation. Nd was the second most abundant metal measured (26.7% Nd). In addition to Nd, the magnet swarf contained significant amounts of the light Ln praseodymium (Pr, 4.35%) and the heavy Ln dysprosium (Dy, 3.34%), both of which have high technological value.[49,50]

To assess the selectivity of *M. extorquens* AM1 uptake for REE, we determined the metal content of the abiotic leachate, and cells and supernatants of cultures grown with 1% (w/v) magnet swarf. Approximately 60% of metal leached by uninoculated Hypho*^MOD^* medium consisted of REE (Figure 2, abiotic control). In cultures grown under the same conditions, most of the available REE was taken up by the cell, leaving the supernatant with 98% Fe (Figure 2). Next, we assessed the specificity of REE bioaccumulation in batch bioreactor cultures with 1% magnet swarf (w/v). Cultures were grown to maximum culture density of 1.8 OD (600 nm) with 1.6 g/L methanol, after which Fe, Nd, Pr, and Dy content in cells and supernatants were measured. Supernatant metal content consisted of mostly Fe (95.9%) with low amounts of REE (Nd, 3.6%; Pr, < 1%; Dy, < 1%) (Figure 4). In cell samples, 98.0% of accumulated metal was REE, 96.8% of which was Nd, while Pr and Dy made up a combined 1.2% (Figure 4). This was ∼1.6-fold higher than what was observed in the 1 mL microplate cultures as mentioned above and could be accounted for by more efficient mixing of swarf with *M. extorquens* AM1 in a stirred-tank system as well as better oxygenation of the culture medium. Fe accounted for only 2.0% of measured intracellular metal content (Figure 4). Given that cells acquire Fe as a micronutrient, it is difficult to differentiate Fe from the growth medium from that obtained from the magnet swarf, but it can be concluded that the latter is less than 2.0%. Together, these data show preferential REE uptake and bioaccumulation in the bioreactor.

**Figure 2.**
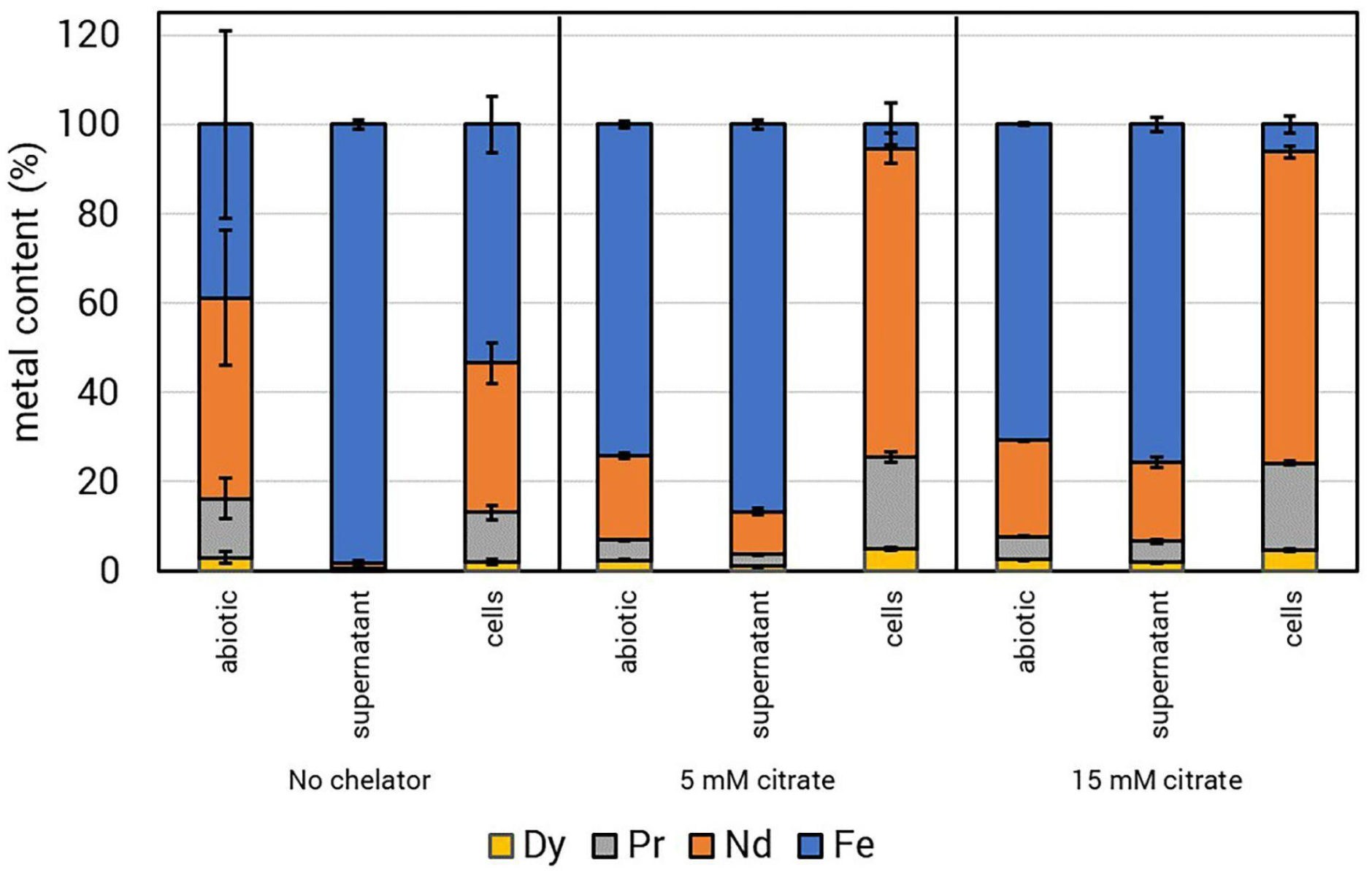
Selective bioconcentration of REE. *M. extorquens* AM1 was grown in 1 mL Hypho*^MOD^*methanol medium with 1% Nd magnet swarf with 0 mM, 5 mM, or 15 mM citrate. Cell and supernatant samples were taken once the culture reached early stationary growth phase. Plots reflect the mean metal contents of three independent growth experiments with error bars showing standard deviations. Dy, dysprosium; Pr, praseodymium; Nd, neodymium; Fe, iron.

### Chemical strategies for increased REE leaching from Nd magnet swarf

Only trace amounts of REE were detected after process runs in bioreactor culture supernatants, indicating that nearly all leached REE were scavenged (Table 1). REE leaching, therefore, could be a limiting factor for bioaccumulation. Microbially-produced organic acids are an environmentally and financially attractive alternative to traditional hydrometallurgical methods.[51–53] Organic acids derived from heterotrophic microbial metabolism have been successfully employed as biolixiviants for metal dissolution, including REEs.[18,19,54–63]

Addition of citric acid to Hypho*^MOD^* medium with magnet swarf increased chemical leaching of all metals (Table S1 and Figure S3). After 28 h of incubation, 8.2 ± 0.1 ppm and 58.7 ± 0.6 ppm of REE were leached into the supernatant supplemented with 5 mM and 15 mM citrate, respectively, which were 15- and 108-fold greater than that found in the supernatant with no chelator (Figure S3). However, Fe was preferentially leached to concentrations of 23.7 ± 2.5 ppm and 142.0 ± 3.2 ppm in the presence of citrate, i.e., there was a 67- and 404-fold increase in Fe concentrations under these conditions (Table S1). The increase in leached Dy with the addition of citrate was twice that of Nd or Pr (Table S1), which may be attributable to its higher complex formation constant with citrate (log*K* = 10.69 vs 9.76 − 9.94).[64] Addition of 55 mM gluconate or 5 mM oxalate, the highest permissible concentration of each for comparable growth of *M. extorquens*, did not increase REE leaching as compared to the negative control condition, though Fe concentrations did increase by 12- and 4.2-fold, respectively (Table S1). Gluconic acid was previously shown to promote leaching of REEs from solid feedstocks, though the pH conditions were more acidic compared to the buffered pH of Hypho*^MOD^*medium (< pH 3.3 vs pH 6.9).[19]

We investigated if nonspecific leaching by the biolixiviant citrate would alter the specificity of REE uptake by *M. extorquens* AM1. When 5 mM or 15 mM citrate was added to the medium, there was a ∼2-fold increase in cellular REE content, from 47% to > 93%, consisting of 69% Nd, 20% Pr, and 5% Dy. The REE content in the culture supernatant also increased from 2% to 13% and 24%, respectively (Figure 2). *M. extorquens* AM1 Δ*mxaF* grew well with 1% (w/v) magnet swarf as the sole source of REE, with significantly higher growth rate as compared to that grown with soluble La (Figures 3A and S4). Addition of citrate to the growth medium with magnet swarf did not significantly affect growth. However, addition of gluconate reduced the growth rate as compared to that of citrate, and oxalate further negatively impacted growth. As expected from the abiotic leaching results (Figure S3), addition of citrate greatly improved REE leaching (Figures 3B and S5A), and led to greater REE uptake by *M. extorquens* AM1 Δ*mxaF* (Figures 3C and S5B). Interestingly, more REE was associated with biomass when 5 mM, rather than 15 mM, citrate was supplemented, suggesting there exists a balance between abiotic leaching and biological uptake at the scale tested (Figure S5B). In fact, upon scale-up of culture volume to 100 mL, 15 mM citrate inhibited growth of *M. extorquens* AM1 Δ*mxaF* (Figure S6). Importantly, unlike the increased uptake of REE in the presence of citrate, the level of Fe associated with the cell remained constant across all conditions (Figure S5B).

**Figure 3.**
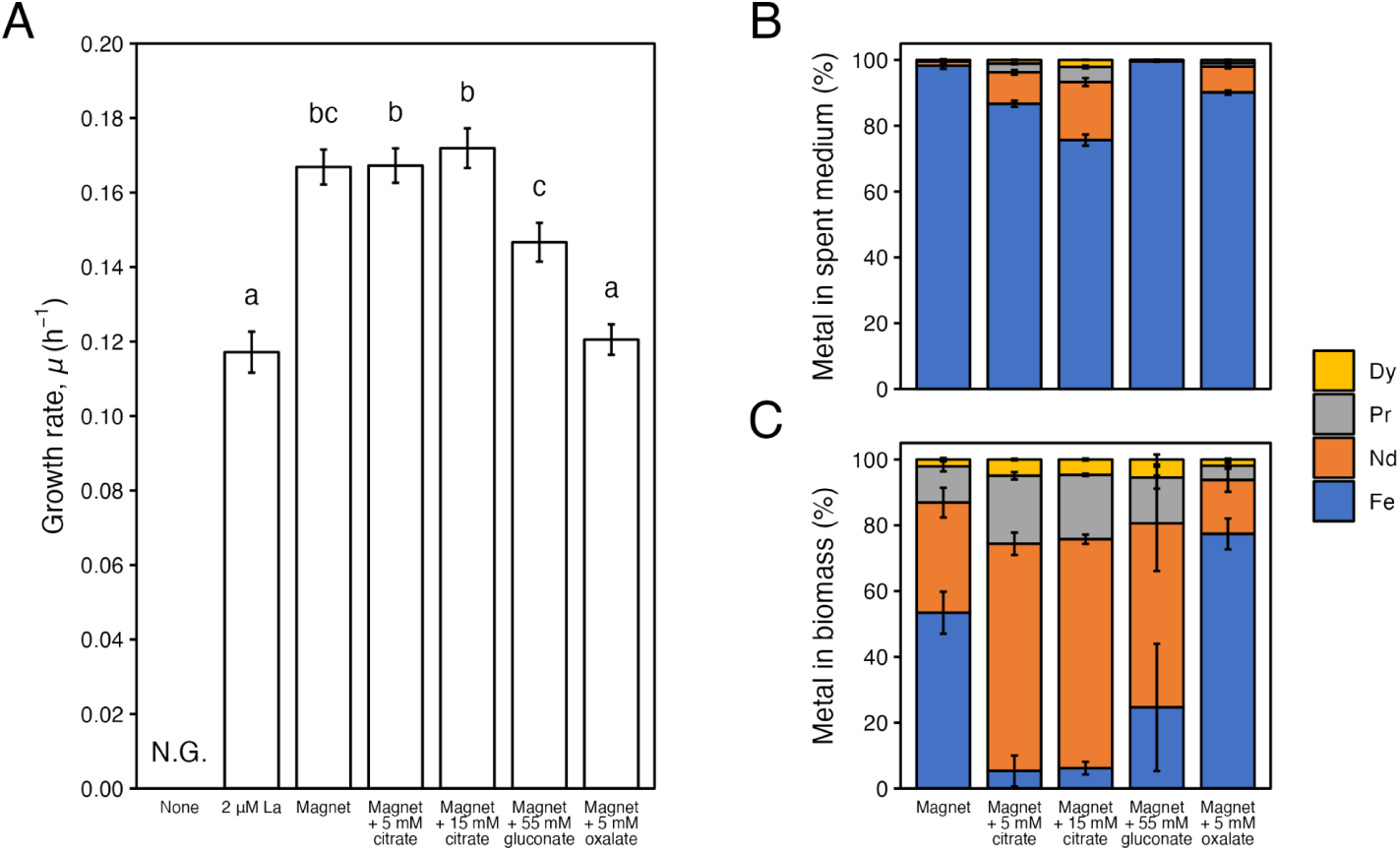
Growth of *M. extorquens* AM1 Δ*mxaF* in Hypho*^MOD^* medium with 1.6 g/L MeOH and 2 μM La, 1% (w/v) magnet swarf, or 1% (w/v) magnet swarf supplemented with citrate (5 mM or 15 mM), gluconate (55 mM), or oxalate (5 mM). (A) Growth rate of *M. extorquens* AM1 Δ*mxaF*, (B) metals leached into the spent medium, and (C) metals associated with biomass were determined based on biological triplicates. Bars and error bars represent model estimate and standard error of growth rates or mean and standard deviation of metal content. Letters above bars represent statistical significance as determined by ANOVA followed by Tukey HSD test (*p* < 0.05). N.G. represents no growth.

Next, we assessed how the addition of 5 mM citrate impacts batch bioreactor cultures and found that total REE bioaccumulation increased to 98.8% of metal uptake with Nd accounting for 88.9%, Pr 9.2%, and Dy 0.7% (Figure 4). Supernatant REE concentrations were notably higher with the addition of citrate to the growth medium. Nd accounted for 24.5%, Pr 2.3%, and Dy 7.9%, with Fe only accounting for 65.2% (Figure 4). Nd and Fe concentrations in citrate-supplemented supernatants were similar to the original swarf composition, but Pr and Dy concentrations were notably different. Pr in the supernatant was ∼2-fold lower and Dy ∼2-fold higher in supernatant, indicating preferential uptake of the light REE and/or preferential leaching of the heavy REE. Overall, these data show highly selective, ∼3.6-fold bioconcentration of REE from magnet swarf by *M. extorquens* AM1. Based on these highly promising results, *M. extorquens* AM1 appears to be naturally poised as a competitive REE bioaccumulation microbe with scalable growth.

**Figure 4.**
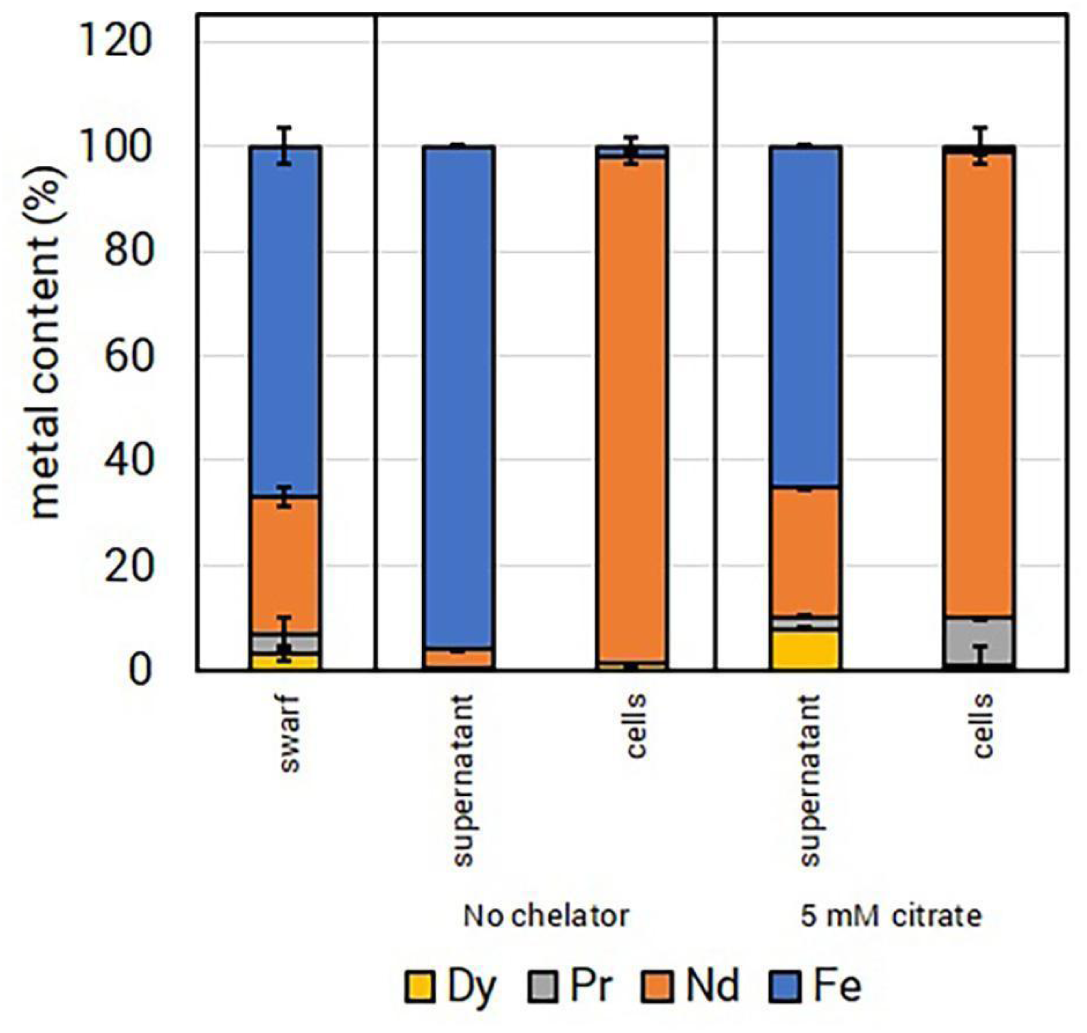
Selective bioconcentration of REE from culture grown in a 0.75 L benchtop bioreactor. *M. extorquens* AM1 was grown in Hypho*^MOD^* methanol medium with 1% Nd magnet swarf. Cell and supernatant samples were taken once the culture reached early stationary growth phase. Metal concentrations in samples were determined by ICP-MS and normalized to the total metal uptake. Metal uptake from cultures with 0 mM or 5 mM citrate added to the growth medium. Plots show the means of three independent growth experiments with error bars showing standard deviations. Dy, dysprosium; Pr, praseodymium; Nd, neodymium; Fe, iron.

Current REE bioaccumulation yields do not account for 100% of the REEin magnet swarf. We tested if a single magnet swarf batch could be processed for further REE extraction in both small- and large-scale cultivations. In 1-mL cultures, the same batch of magnet swarf was able to sustain growth of *M. extorquens* AM1 Δ*mxaF* for five cycles, though the final OD and total leached REE concentrations decreased with continuing growth cycles (Figure S7). Based on the bioaccumulation yields from the first bioreactor run (Figure 4), we estimated 8.25 ± 0.04% of the swarf Nd could be recovered in a single fed-batch process. With a second consecutive process run, using the same swarf batch as the first run, bioaccumulation of Nd accounted for another 7.75 ± 0.12% of the original amount (Figure 5). These promising results suggest that a continuous process run could feasibly be used to recover an even higher proportion of REE in a swarf batch.

**Figure 5.**
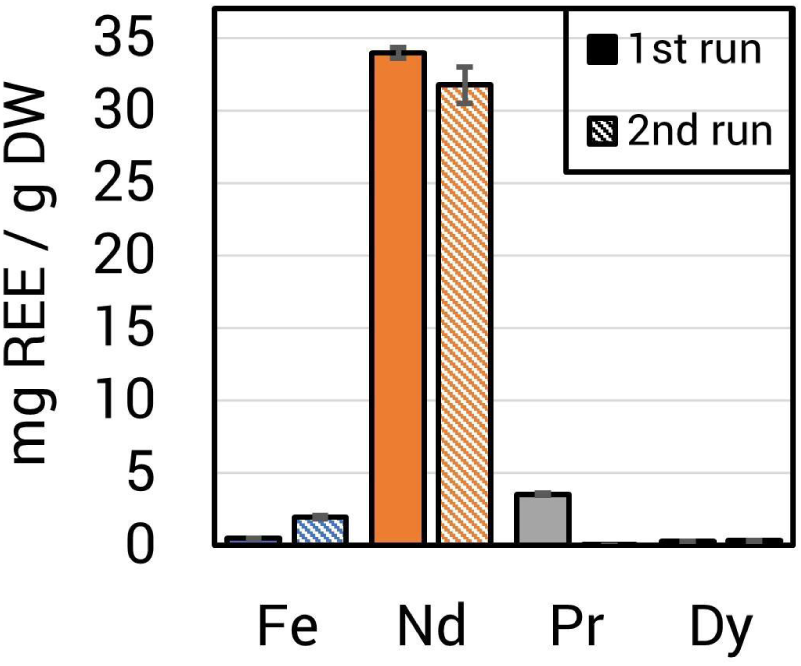
Total REE bioaccumulation is increased by reusing magnet swarf feedstock. Each experiment consisted of two complete process runs in which the Δ*mxaF* strain was grown with 1% magnet swarf (w/v) in a 0.75 L bioreactor with 1.6 g/L methanol and 5 mM citrate. After growth, cell samples were taken for ICP-MS metal content measurements, and the magnet swarf was removed from the bioreactor, rinsed 4 times with ddH_2_O, and dried at 65°C for 72 h before reuse. Bars show the means of cellular REE content normalized to dry weight (DW) for two replicate experiments. A distinct swarf batch was used for each individual experiment. Error bars indicate standard deviations.

A recent techno-economic analysis reported the potential economic feasibility of using biolixiviants to increase REE leaching from waste materials and found that the carbon substrate (glucose) is a major contributor to cost.[20] Our results showed that addition of low molarity citrate significantly increased REE leaching and bioaccumulation from Nd magnet swarf. Citrate can be added directly to the bioreactor, or in future process development could be linked via co-culture with a heterotrophic acid-producing microbe, such as *Gluconobacter oxydans*. However, to truly limit the costs of an REE recovery process, an acid-independent approach would be preferred, especially in areas where secondary hazardous waste streams are of concern.

### Approaches for enhanced REE-specific bioleaching and bioaccumulation

The genetic tractability and availability of genetic tools for strain engineering make *M. extorquens* AM1 a uniquely suitable microbial platform for REE recovery. A biosynthetic pathway for an REE-chelator (lanthanophore) has been previously reported and was named the LCC (lanthanide <u>c</u>helation <u>c</u>luster), according to the function of its predicted product.[65] Expression of the LCC biosynthetic product has already been shown to increase Nd uptake in *M. extorquens* AM1 from both soluble chloride and poorly soluble oxide forms in a highly specific manner.[65] We tested the impact of expressing the LCC in trans during growth with magnet swarf and saw increases in REE bioaccumulation. Nd, Pr, and Dy accumulation increased by more than 3-fold with LCC expression, reaching 80 mg Nd/g DW, 15 mg Pr/g DW, and 8 mg Dy/g DW (Figure 6A), while Fe accumulation increased only marginally. Notably, expression of the LCC was under control of the *lac* promoter, a comparatively weak promoter in *M. extorquens* AM1.[66] Generation of a stronger expression system is underway, and in theory should boost REE bioaccumulation even further.

**Figure 6.**
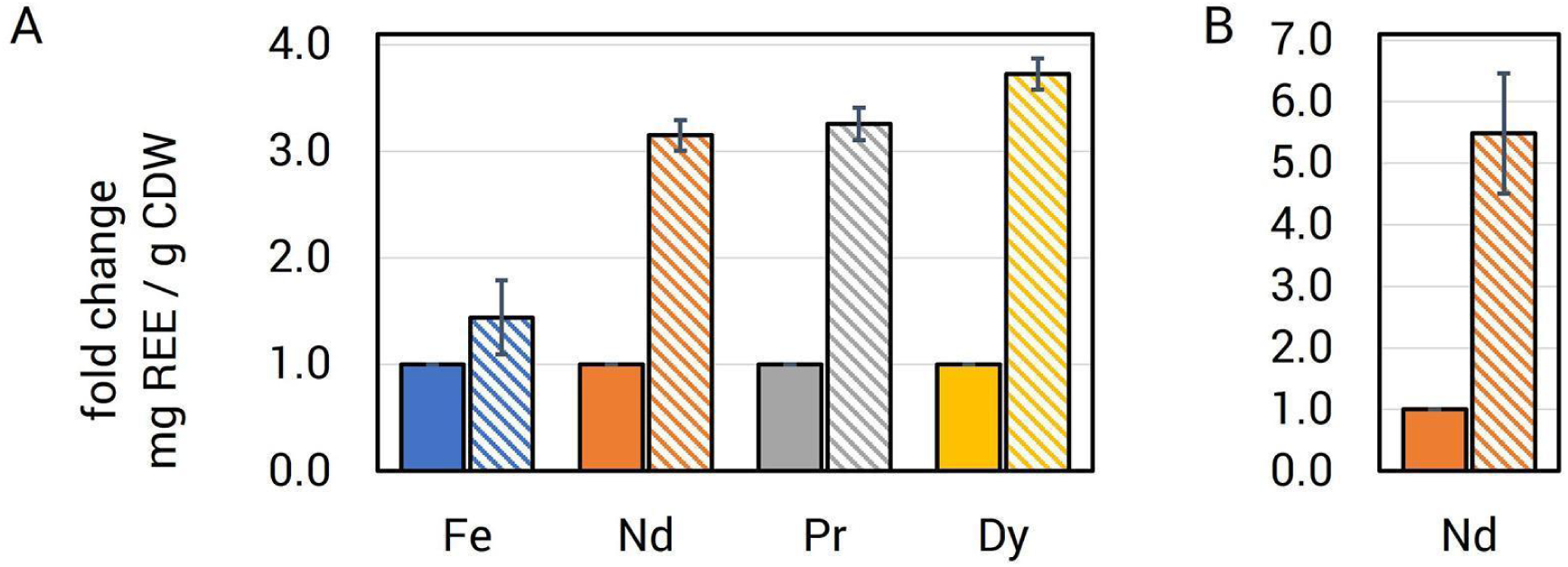
Strain engineering for increased REE bioaccumulation. *A*, in trans expression of the lanthanophore produced from the LCC enhances REE (Nd, neodymium; Pr, praseodymium; Dy, dysprosium) bioaccumulation from magnet swarf in *M. extorquens* AM1, with minimal Fe uptake. Δ*mxaF*, solid bars; Δ*mxaF*/pLCC, hatched bars. *B*, deletion of the exopolyphosphatase encoding *ppx* gene results in higher Nd bioaccumulation. Δ*mxaF*, solid bars; Δ*ppx*, hatched bars. For *A* and *B,* metal content was determined by ICP-MS and normalized to cell dry weight. Values represent the mean of samples from 3 independent cultures grown in shake flasks. Error bars reflect standard deviations.

Pyrroloquinoline quinone (PQQ) has been shown to directly bind REEs [67] and has been implicated in microbial REE solubilization.[63] We recently reported *M. extorquens* AM1 strain evo-HLn[40] that exhibited enhanced PQQ production in the presence of the heavy REE, gadolinium. We measured Nd bioaccumulation in evo-HLn and saw a 53% increase compared to Δ*mxaF*.

REEs are stored in intracellular polyphosphate granules in *M. extorquens* AM1. Depolymerization of polyphosphate is catalyzed by exopolyphosphatase activity. We hypothesized that limiting the cellular capacity for polyphosphate depolymerization could generate higher levels of REE bioaccumulation. A *ppx* (encoding exopolyphosphatase) deletion strain was generated and assessed for REE bioaccumulation, showing a ∼5.5-fold increase in Nd levels reaching 202 mg Nd/g DW (Figure 6B).

The results shown herein demonstrate that selective targeting of specific processes for REE metabolism, generated by a detailed understanding of the physiology of *M. extorquens* AM1, is an effective strategy for engineering enhanced REE storage. Reduction of inorganic phosphate in the growth medium, likely allowing for increased uptake and granulation with REE, resulted in a nearly 4-fold enhancement of Nd yields over baseline levels (Figure 7). Identification of REE-binding ligands like the LCC biosynthetic product allows for non-acidic, highly specific leaching from feedstocks. Genetic engineering to increase the LCC product resulted in Nd yields of more than 20-fold higher than baseline values (Figure 7). Likewise, detailed knowledge of REE storage granulation and storage processes has allowed for genetic manipulation and engineering to further boost intracellular REE content. By engineering reduced capacity for granule deconstruction, we were able to increase Nd storage capacity over 50-fold (Figure 7) relative to the baseline condition with Δ*mxaF*. Our current yields when reaching an OD of 20 in bioreactor would allow Nd recovery of 1.3 g - 2.1 g of Nd/L representing a recovery between 65%-100% of Nd in a single process run when using 1% Nd magnet swarf pulp density.

**Figure 7.**
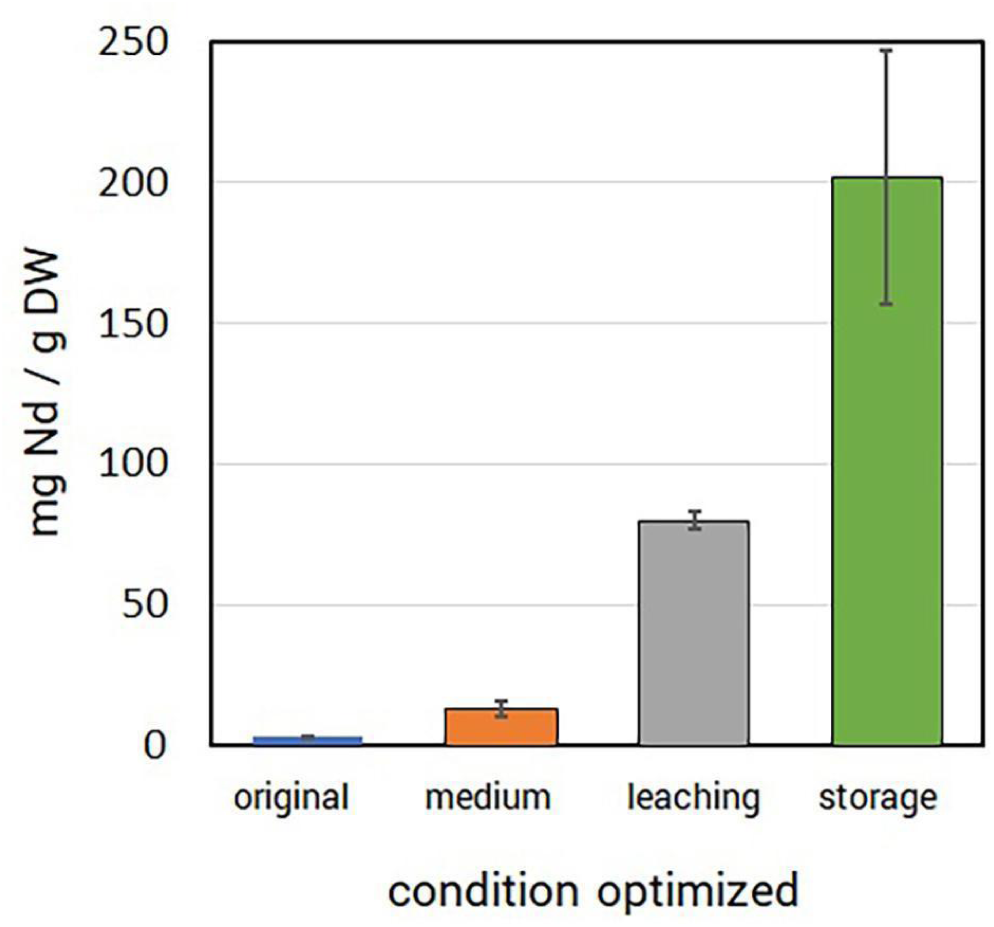
Process performance enhancement. Chemical and genetic targeting of specific metabolic processes generates improved REE bioaccumulation relative to original (baseline) measurements. Manipulations to growth medium inorganic phosphate (medium), lanthanophore production (leaching), and REE/phosphate granulation (storage) each generated elevated REE bioaccumulation yields. Bioaccumulation was measured by ICP-MS and is normalized to dry weight. Error bars represent standard deviations of biological triplicates.

## CONCLUSIONS

Current REE mining and refinement practices are costly, environmentally destructive, and unsustainable.[68] Safer, cleaner alternatives are needed. Microbes have emerged as a promising biological alternative to harsh chemical extraction methods for leaching REE from complex sources and waste streams, opening new avenues for processing feedstocks and recycling these critical metals. *M. extorquens* AM1, a bacterium with the natural ability to solubilize and accumulate REE, provides a unique pathway towards selective, REE bioleaching, bioaccumulation, and recovery that is scalable and does not rely on extremophile organisms and their inherent limitations. Future work will investigate this further with a focus on end-to-end conversion of feedstock to salable rare earth oxide products.

We have reported optimization of several parameters to increase REE recovery yields in a 10 L bioreactor format reaching yields of over 1.3% dry weight, rivaling current state of the art technologies. The potential *M. extorquens* AM1 to recover REE is not limited to E-waste, as our process is compatible with even the most unrefined sources, such as monazite and bastnäsite ores, and pulverized smartphones (Table S2). *M. extorquens* AM1 is uniquely suited biologically for REE recovery through its production of lanthanophores, dedicated REE transport machinery, and its capability for producing REE-phosphate granules that will facilitate purification. The potential reduction in REE-mining and -refinement pollution and hazardous exposure make *M. extorquens* AM1 an attractive, green alternative to current chemical methodologies.

## Author Contributions

The manuscript was written through contributions of all authors. All authors have given approval to the final version of the manuscript.

## Funding Sources

The information, data, or work presented herein was funded in part by the Advanced Research Projects Agency-Energy, United States Department of Energy, under Award Number DE-AR0001337.

## ACKNOWLEDGMENTS

We would like to thank Yi Yao and Samir Budhathoki from the University of Wyoming for their assistance with some ICP-MS sample analysis, and Bang Luong, Caitlin Hoeber, Clarisse Hufana, Deval Patel, James Cai, Kim Luu, Krisha Gupta, Mariam Jacob, Nicholas Lien, Alice Huang, and Jennifer Lepe-Rodriguez for assistance with CFU measurements for smartphone growth curves. Part of this work was carried out under the auspices of the DOE by Lawrence Livermore National Laboratory under contract DEAC52-07NA27344 (LLNL-JRNL-851205).

## SUPPORTING INFORMATION

Supplemental methods; Figure S1, Impact of inorganic phosphate on strain performance; Figure S2, Effect of calcium in growth medium on Nd uptake; Figure S3, Abiotic leaching of REE; Figure S4, Growth rates of *M. extorquens* AM1 with organic acid addition; Figure S5, bioleaching and bioaccumulation measurements after growth with organic acids; Figure S6, Effect of citrate on bioleaching and bioaccumulation; Figure S7, Iterative bioaccumulation from a single swarf batch; Table S1, Ratio of metals chemically leached from magnet swarf by organic acids; Table S2, Growth of *M. extorquens* AM1 with pure and complex REE sources.

## TOC graphic

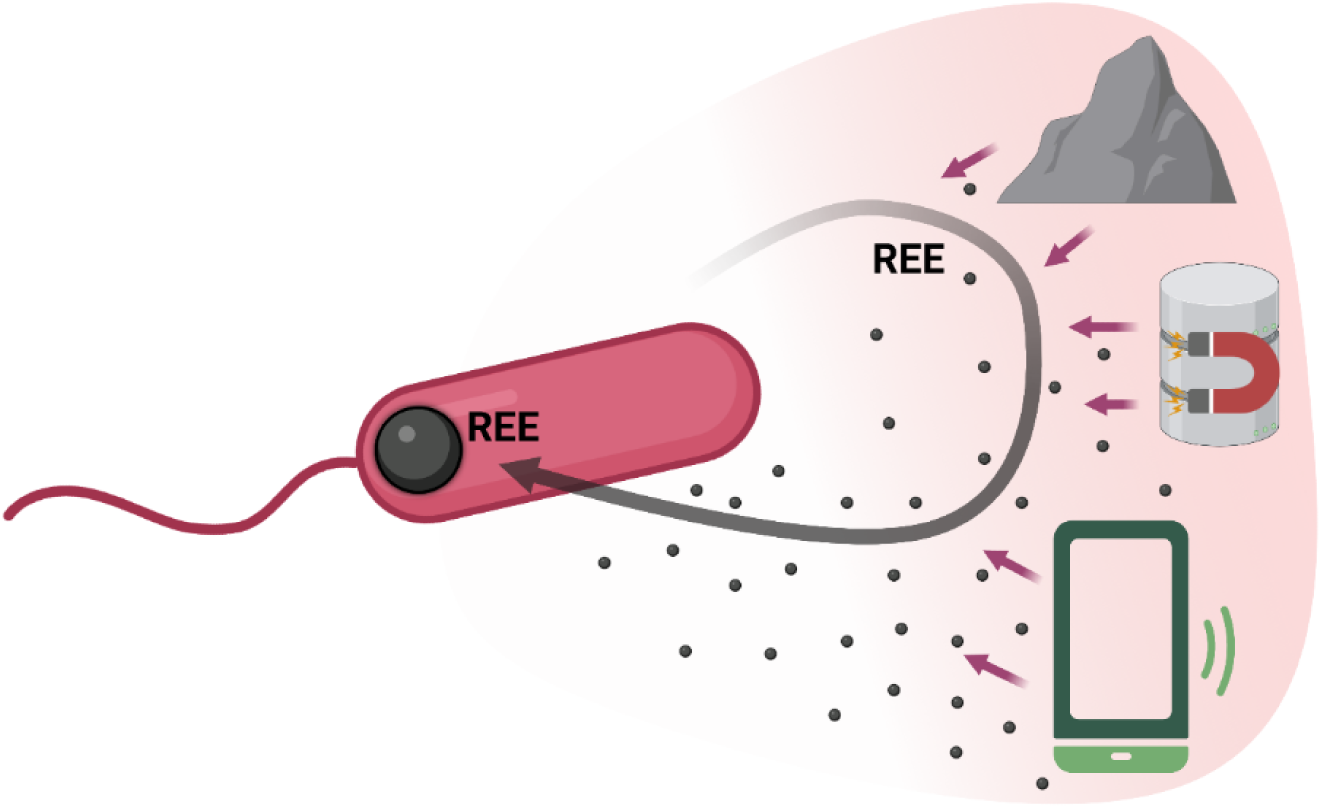

**For table of contents only**

## SYNOPSIS

REE production requires harsh acids, high temperatures and pressures, is costly, and is environmentally taxing. This study shows a scalable and sustainable bacterial platform for non-acidic, targeted recovery of REE from E-waste.

## Notes

### Competing Interest Statement

The authors have declared no competing interest.

