## Supporting Material for "Scalable bio-platform to recover critical metals from complex waste sources"

**Number of Pages: 14**

**Number of Figures: 7**

**Number of Tables: 1**

*SUPPLEMENTAL METHODS*

**Hypho medium optimization.** *M. extorquens* AM1 pre-cultures were grown as described. Cells were pelleted by centrifugation at 1,000 x *g* for 10 minutes, washed in 1 mL sterile Hypho medium, and then resuspended in 200 µL sterile Hypho medium. 48-well tissue culture plates (Corning, New York) were prepared with 640 µL of medium prepared with standard phosphate ($P_i$; 2.53 g/L $K_2HPO_4$, 2.59 g/L $NaH_2PO_4.H_2O$), half phosphate (½ $P_i$; 1.27 g/L $K_2HPO_4$, 1.30 g/L $NaH_2PO_4.H_2O$; Hypho$^{MOD}$ medium), or quarter phosphate (¼ $P_i$; 0.63 g/L $K_2HPO_4$, 0.65 g/L $NaH_2PO_4.H_2O$) and inoculated with 10 µL of cell suspension. Cultures were grown in an Epoch II microplate reader (BioTek, Winooski, VT) for 48 hours, shaking at 548 rpm, incubated at 30ºC. Growth of cultures was monitored by measuring light scatter at 600 nm every 30 minutes.

**Processing of smartphones.** Postconsumer ZTE Quest N817 and Nokia 6136 smartphones including lithium ion batteries were purchased from eBay. Phones were pulverized using a Blendtec Total Blender (Blendtec, Orum, UT) in pulses of ~20 seconds for 5 to 10 min. The resulting debris was sifted through a set of geological sieves (American Geo, Poughkeepsie, NY). Fragments below 0.15 mm were retained and autoclaved.

**Growth analysis with ores, minerals, and smartphones.** For growth with ores and minerals, cultures of *M. extorquens* AM1 were grown overnight in MP succinate medium without exogenous lanthanides. When cultures reached mid-log phase (~OD 0.8), a 300 µL aliquot was sub-cultured to 10 mL of fresh MP methanol supplemented with 0.5% ore. As controls, 10 mL MP methanol with ores and without or with the addition of 2 µM lanthanum chloride ($LaCl_3$) was inoculated with 300 µL bacterial aliquots or with MP methanol lacking cells. Every 3 hours, a 200 µL aliquot of culture was removed and $OD_{600}$ was recorded using a Spectramax M2 plate reader (Molecular Devices, Sunnyvale, CA). The no cell control containing the respective ore was used as the blank. For growth with smartphones, 2 mL of MP succinate culture was inoculated into 100 mL MP methanol media with and without 0.5% smartphone powder and $LaCl_3$ in 250 mL shake flasks. Growth was measured by serial dilution and CFU analysis.

*SUPPORTING FIGURES*

**Inorganic phosphate optimization in growth medium.** REEs are poorly soluble as phosphate compounds,[1] the formation of which could hinder REE-dependent methanol growth in media with high phosphate concentrations. Our initial studies showed that reducing inorganic phosphate in the growth medium resulted in better growth performance (Figure S1). For scale-up growth experiments we therefore chose to use Hypho$^{MOD}$ medium containing half the standard phosphate concentration (see Methods).

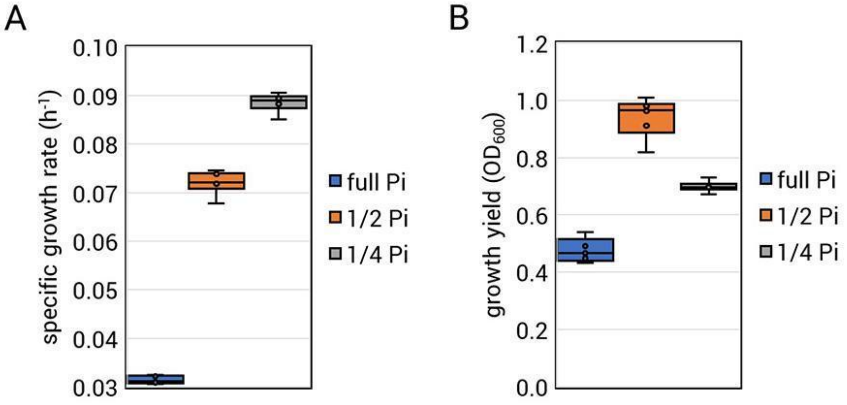


**Figure S1.** Impact of inorganic phosphate on strain performance. *A,* Growth rate of *M. extorquens* AM1 increases with decreasing inorganic phosphate in the medium. Wild type *M. extorquens* AM1 was grown in a 48-well microplate in Hypho minimal medium with 1.6 g/L methanol and 0.5 mg/L $NdCl_3$. Blue, standard phosphate ($P_i$; 2.53 g/L $K_2HPO_4$, 2.59 g/L $NaH_2PO_4.H_2O$). Orange, half phosphate (½ $P_i$; 1.27 g/L $K_2HPO_4$, 1.30 g/L $NaH_2PO_4.H_2O$; Hypho$^{MOD}$ medium). Gray, quarter phosphate (¼ $P_i$; 0.63 g/L $K_2HPO_4$, 0.65 g/L $NaH_2PO_4.H_2O$). *B*, Yield of *M. extorquens* AM1 grown on methanol with Nd is influenced by inorganic phosphate in the medium. Box plots represent the interquartile ranges for 6 biological replicates; standard deviation is shown by error bars.

Calcium is normally added to Hypho minimal media as an essential metal for MxaF methanol dehydrogenase. However, the strains of *M. extorquens* AM1 used in this study do not have *mxaF* and rely on REE for methanol dehydrogenase activity via XoxF and ExaF. We tested if removal of calcium from the growth medium would impact REE uptake in *M. extorquens* AM1 but observed no significant differerences (Figure S2).

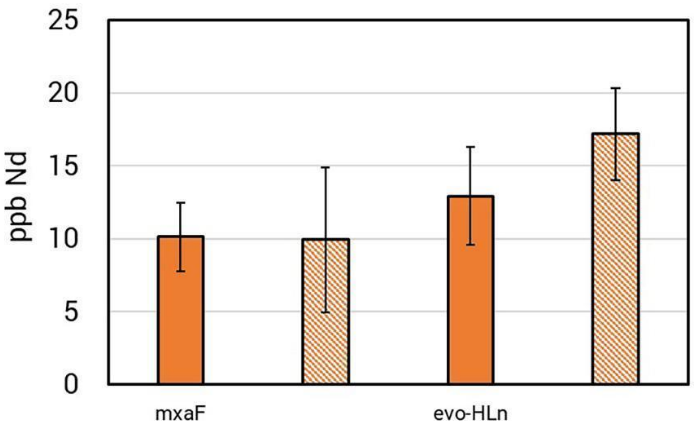


**Figure S2.** Effect of calcium in growth medium on Nd uptake. *M. extorquens* AM1 Δ*mxaF* and evo-HLn strains were grown in Hypho$^{MOD}$ medium with 1.6 g/L methanol, with (solid bars) or without (hatched bars) 2 μM $CaCl_2$, to early stationary phase. Nd content was determined by ICP-MS and normalized by OD. Bars show the means of three biological replicates; standard deviation is shown by error bars. No significant differences were observed among conditions by Student's T-test.

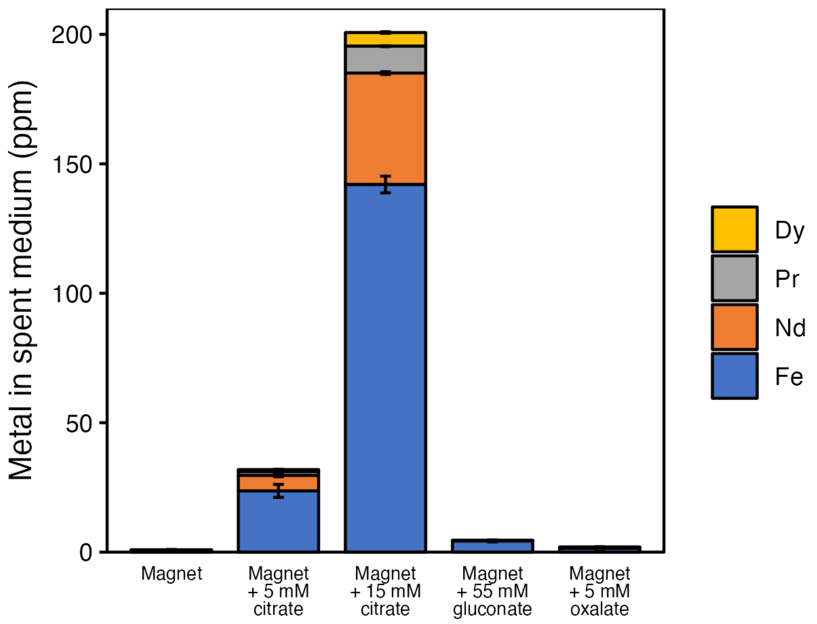


**Figure S3.** Abiotic leaching of REE from 1% (w/v) magnet swarf in 1 mL Hypho$^{MOD}$ medium supplemented with 5 mM or 15 mM citrate, 55 mM gluconate, or 5 mM oxalate, incubated at 30 °C, as determined by ICP-MS. Each bar and error bars represent mean and standard deviation of triplicate samples, respectively.

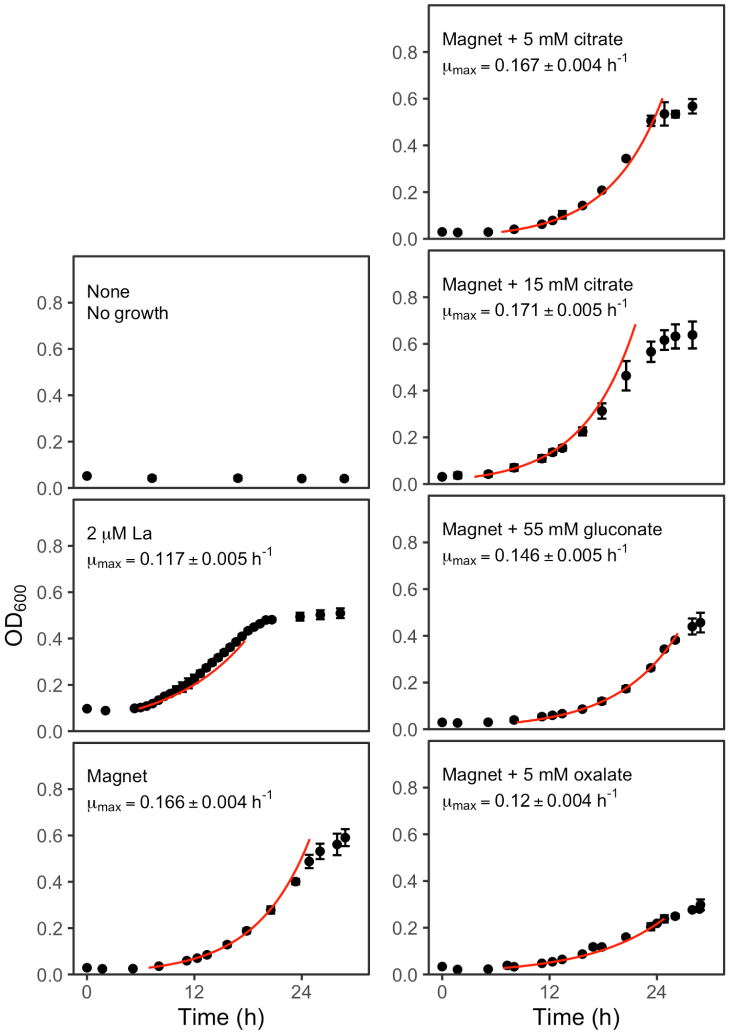


**Figure S4.** Growth rates of *M. extorquens* AM1 Δ*mxaF* in Hypho$^{MOD}$ medium with 1.6 g/L MeOH, supplemented with La or 1% (w/v) magnet swarf as source of REE and various organic acids. Each point and error bar represent mean and standard deviation of biological triplicates. Red line represents model fit as determined using the R package growth rates on at least three biological replicates. Growth experiments were performed in 1-mL volume in 24-well plates.

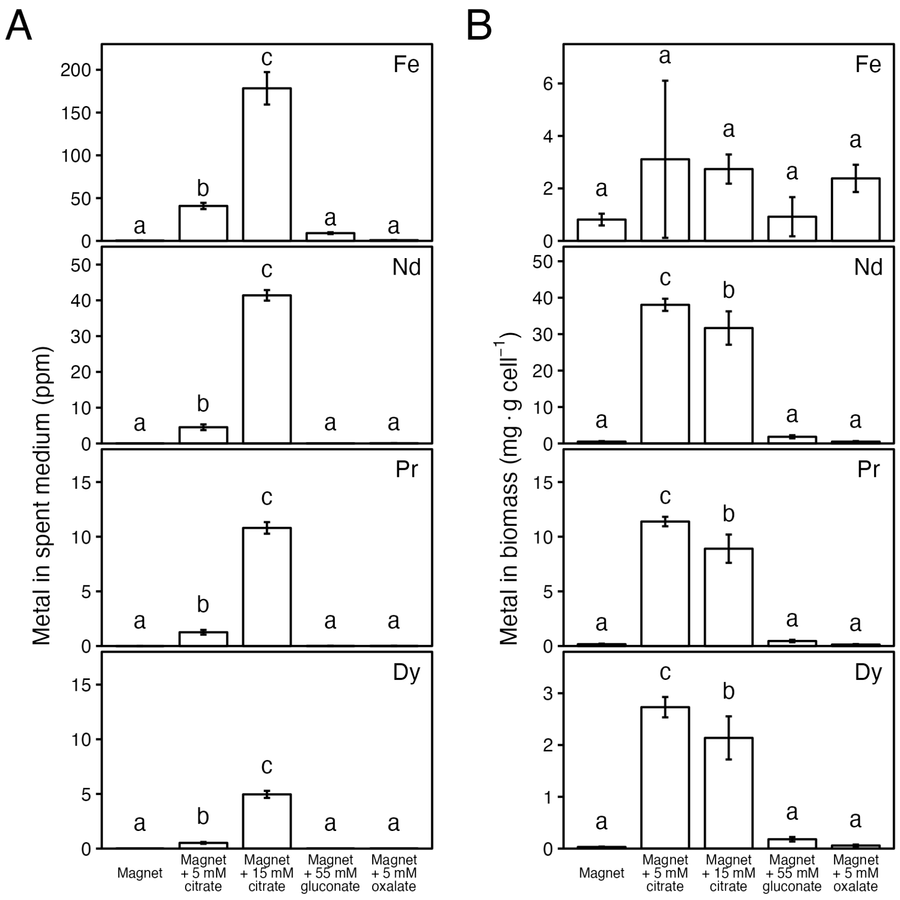


**Figure S5.** Bioleaching and bioaccumulation during growth with organic acids. Amount of Fe, Nd, Pr, and Dy found in (A) the spent medium and (B) biomass of *M. extorquens* AM1 Δ*mxaF* after growth in 1 mL Hypho$^{MOD}$ with 1.6 g/L MeOH and 1% (w/v) magnet swarf, supplemented with 5 mM or 15 mM citrate, 55 mM gluconate, or 5 mM oxalate. Metal concentrations were determined by ICP-MS. Bars and error bars represent mean and standard deviation of triplicate samples. Letters above bars represent groups of statistical significance as determined by ANOVA followed by Tukey HSD test ($p < 0.05$).

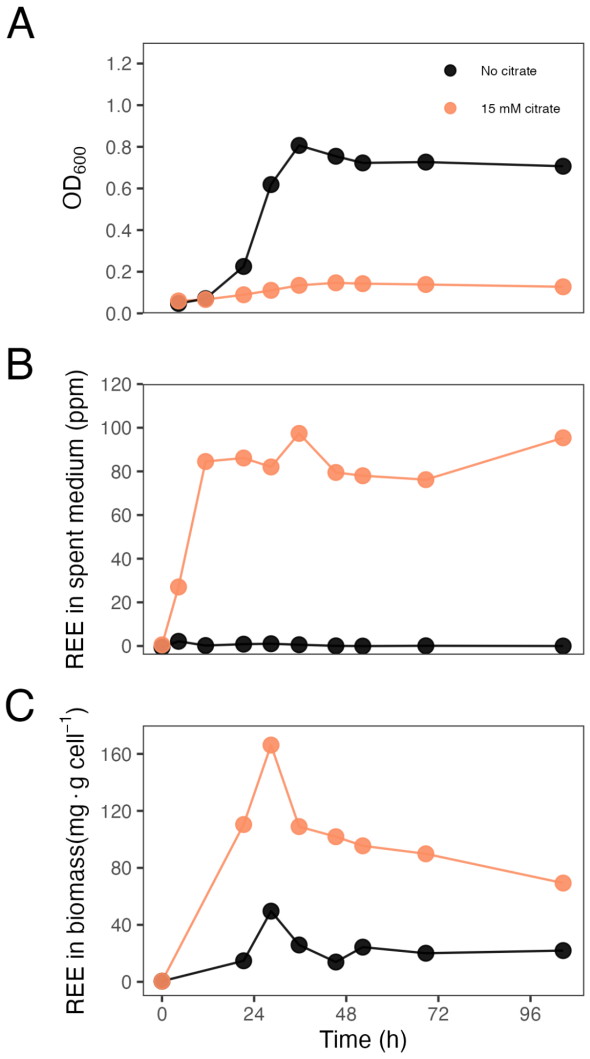


**Figure S6.** Effect of citrate on bioleaching and bioaccumulation. (A) Growth of *M. extorquens* AM1 Δ*mxaF* in 100 mL of Hypho$^{MOD}$ medium with 1.6 g/L MeOH, with and without 15 mM citrate. Concentrations of REE in (B) spent medium and (C) cell pellets were analyzed using the Arsenazo III assay. Each point represents one biological replicate.

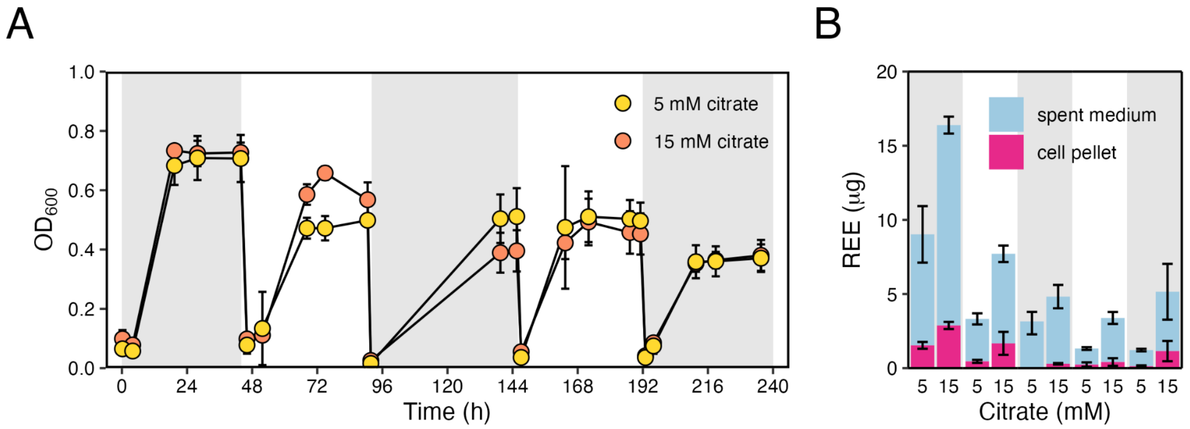


**Figure S7.** Iterative bioaccumulation from a single swarf batch. Growth of *M. extorquens* AM1 Δ*mxaF* in 1 mL of Hypho$^{MOD}$ medium with 1.6 g/L MeOH, 1% (w/v) magnet swarf, and varying concentrations of citrate. (A) Five cycles of growth were repeated for the same magnet swarf sample by replacing culture with fresh medium, then (B) REE in spent medium and cell pellet analyzed via Arsenazo III assay. Each point or bar represents the mean of biological triplicates, and error bars the standard deviation.

*SUPPORTING TABLES*

**Table S1.** Ratio of metals chemically leached from magnet swarf in the presence of citrate, gluconate, and oxalate as compared to the absence of chelator (mean ± SD).

| | Fe | Nd | Pr | Dy |
|---|---|---|---|---|
| Magnet (reference) | 1.0 ± 0.6 | 1.1 ± 0.3 | 1.0 ± 0.3 | 1.2 ± 0.5 |
| Magnet + 5 mM citrate | 67.4 ± 7.2 | 15.8 ± 1.4 | 13.0 ± 0.9 | 33.5 ± 5.1 |
| Magnet + 15 mM citrate | 404.4 ± 9.2 | 113.8 ± 1.4 | 92.3 ± 1.5 | 223.0 ± 13.6 |
| Magnet + 55 mM gluconate | 12.1 ± 0.9 | 0.6 ± 0.2 | 0.6 ± 0.2 | 0.7 ± 0.3 |
| Magnet + 5 mM oxalate | 4.2 ± 0.5 | 1.0 ± 0.4 | 1.0 ± 0.4 | 1.6 ± 0.6 |

**Table S2.** Growth of *M. extorquens* AM1 with pure and complex REE sources

| REE source | growth rate |
|---|---|
| 2 μM $LaCl_3$ | 0.16 ± 0.01 |
| 0.5% monazite ore | 0.15 ± 0.03 |
| 0.5% bastnasite ore | 0.16 ± 0.00 |
| 0.5% bastnasite crystal | 0.16 ± 0.01 |
| 0.5% blended smartphone | 0.16 ± 0.01[є] |

[є]growth rate determined from cfu measurements every 4 h.

***Flexibility of M. extorquens for REE bioleaching and bioaccumulation from complex sources***

Robust growth and bioaccumulation of Nd from magnet swarf indicated that *M. extorquens* AM1 can effectively leach and acquire Ln from poorly soluble sources. Therefore, we tested the ability of *M. extorquens* AM1 to grow with unrefined REE sources with very low solubility in the growth medium. Monazite ore, for example, contains up to ~45% REEs, with the $Ce_2O_3$ content comprising as much as 17% of the total REEs. However, the REEs occur as poorly soluble phosphates and fluorocarbonates in these ores. When challenged with the ores monazite or bastnäsite as the sole REE source, *M. extorquens* AM1 grew as well as with soluble $LaCl_3$ (Table 2). Similar growth results were also observed with bastnäsite crystal (Table S2).

Complex debris like E-waste represents virtually untapped reservoirs of valuable REEs. Cellular smartphones contain several species of REE, including yttrium, lanthanum, terbium, neodymium, gadolinium and praseodymium, making smartphone E-waste a valuable, unutilized source for REE recovery. However, smartphone E-waste poses two major challenges for bioleaching. The metals in this E-waste are highly insoluble in oxide form, and smartphone batteries contain other metals (*e.g.,* iron, boron, copper, tellurium, manganese, mercury, lead, tungsten, lithium, cobalt), all of which can be toxic. We tested the feasibility of using smartphone E-waste as a REE source by assessing the capacity of *M. extorquens* AM1 to grow on methanol. Somewhat surprisingly, with 0.5% pulp density (w/v) of blended smartphones, *M. extorquens* AM1 grew as well as with soluble $LaCl_3$ (Table S2). In fact, along with the ores and crystal REE sources, the insolubility of the metals and complexity of the source did not significantly impact growth rate or growth yield of the culture (Table S2) showing the flexibility and robustness of *M. extorquens* AM1 as a biological platform for the bioaccumulation of REE from complex sources.

*REFERENCES*

(1) Firsching, F. H.; Brune, S. N. Solubility products of the trivalent rare-earth phosphates. *J. Chem. Eng. Data* **1991**, *36* (1), 93–95. https://doi.org/10.1021/je00001a028.
